# An Efficient Direct Conversion Strategy to Generate Functional Astrocytes from Human Adult Fibroblasts

**DOI:** 10.1101/2024.09.02.610876

**Authors:** Uchit Bhaskar, Emily Shrimpton, Jason Ayo, Asiya Prasla, Mark Z Kos, Melanie A. Carless

## Abstract

Direct reprogramming approaches offer an attractive alternative to stem-cell-derived models, allowing the retention of epigenetic information and age-associated cellular phenotypes, and providing an expedited method to generate target cell types. Several groups have previously generated multiple neuronal subtypes, neural progenitor cells, oligodendrocytes, and other cell types directly from fibroblasts. However, while some groups have had success at the efficient conversion of *embryonic* fibroblasts to astrocytes, they have not yet achieved similar conversion efficiency for *adult* human fibroblasts. To generate astrocytes for the study of adult-stage disorders, we developed an improved direct conversion strategy employing a combination of small molecules to activate specific pathways that induce trans-differentiation of human adult fibroblasts to astrocytes. We demonstrate that this method produces mature GFAP+/S100β+ cells at high efficiency (40-45%), comparable to previous studies utilizing embryonic fibroblasts. Further, <u>F</u>ibroblast-<u>d</u>erived induced <u>A</u>strocytes (FdiAs) are enriched for markers of astrocyte functionality, including ion-channel buffering, gap-junction communication, and glutamate uptake; and exhibit astrocyte-like calcium signaling and neuroinflammatory phenotypes. RNA-Seq analysis indicates a close correlation to human brain astrocytes and iPSC-derived astrocyte models. Fibroblast-derived induced astrocytes provide a useful tool in studying the adult brain and complement existing *in vitro* models of induced neurons (iNs), providing an additional platform to study adult-stage brain disorders.

## Introduction

In modeling age-related neurological disorders, cellular reprogramming approaches generating neurons directly from fibroblasts or other somatic cell types are considered to be widely successful (Huh et al., 2016; Koch, 2015; Mertens et al., 2015). In particular, direct conversion strategies avoid a pluripotent stem cell state that is known to erase host epigenetic hallmarks (Horvath, 2013; Mertens et al., 2015, 2021), preserve individual genetic mosaicism, retain cellular aging features, and offer a quicker alternative to iPSC-based reprogramming (Mertens et al., 2018). However, while such models are now routinely used in generating neuronal cell types, similar approaches to generating other brain cell types have been limited.

A growing body of evidence indicates that astrocytes, long believed to be the major supporting cell type in the CNS, have essential functions for normal brain physiology including phagocytosis, glutamate regulation, neuronal and synaptic maturation, trophic support, blood-brain barrier formation and neuroinflammation (Eroglu and Barres, 2010; Verkhratsky and Butt, 2007). It is therefore not surprising that these cells contribute to many brain disorders. iPSC-derived models (Tcw et al., 2017; Voulgaris et al., 2022) have been used to understand astrocyte functionality in healthy and diseased states. However, like neuronal models, stem cell-derived astrocyte models are likely more representative of fetal brain, and direct conversion approaches could alleviate some of these concerns. However, to date, only three studies have reported successful conversion of fibroblasts directly to induced astrocytes (iAs) using either lentiviral expression of transcription factors or a combination of small molecules (Caiazzo et al., 2015; Quist et al., 2022; Tian et al., 2016). Although these methods show high efficiency for embryonic fibroblasts (2-40%), neither approach has been able to efficiently convert human *adult* fibroblasts to functional astrocytes. Tian et. al. observed around 8-10% efficiency (assessed by % GFAP/S100β positive cells) in converting human adult fibroblasts (71-year-old male) to iAs using small molecules (Tian et al., 2016), while Quist et. al. reported that their lentiviral-mediated conversion of postnatal human fibroblasts was not as efficient as in embryonic fibroblasts (Quist et al., 2022).

In this study, we employed a small-molecule cocktail in a three-step conversion process that shows a substantially improved method to generate human adult <u>F</u>ibroblast-<u>d</u>erived induced-<u>A</u>strocytes (**FdiAs**). Utilization of small molecules has been thought to overcome poor induction efficiency and genomic integration of viral vectors, thus offering a safer, easier alternative (Li et al., 2015; Tian et al., 2016). Our method yields FdiAs that exhibit mature phenotypic and transcriptional properties of adult astrocytes and exhibit astrocyte-relevant functionality. In addition, we show that this method is applicable to fibroblasts derived from individuals of different ages and sex, thus providing a powerful *in vitro* model system for the study of brain disorders.

## Results

### A cocktail of small molecules can directly convert human adult fibroblasts to induced astrocytes

Existing lentivirus-mediated direct conversion protocols to generate FdiAs employ transcriptional activation of either a combination of *NFIA/NFIB/SOX9* (Caiazzo et al., 2015) or just *NFIB/SOX9* (Quist et al., 2022) to induce a gliogenic switch. However, previous studies have shown that *NFIA* activation occurs downstream of *SOX9* and that the SOX9-NFIA complex regulates a subset of genes involved in gliogenesis (Kang et al., 2012). SOX9 is believed to be a master regulator of glial fate choice, preventing neurogenesis (Stolt et al., 2003), and prior research in chondrogenic studies indicates that *SOX9* expression could be enhanced by small molecules including fibroblast growth factor 2 (FGF2) (Handorf and Li, 2011) and epidermal growth factor 1 (EGF1) (Vinukonda et al., 2016), in conjunction with bone morphogenic protein 2 (BMP2) (Pan et al., 2008). Further, as evidenced by Tian et.al., TGF-β and GSK-3β inhibition are both critical contributors to the conversion process (Tian et al., 2016). We emulated this by the addition of SB-431542, A-83-01 and CHIR99021 during induction. cAMP, important in signal transduction pathways, is known to promote differentiation and astrocyte development (McManus et al., 1999; Paco et al., 2016). We used a cAMP-activator, forskolin (FSK; during induction), or db-cAMP (during differentiation and maturation) to initiate this process. iPSC-derived astrocyte (**iPSC-iA**) studies have established a key role for BMP4 and JAK-STAT signaling pathways in enhancing GFAP expression, which allows astrocyte development (Krencik et al., 2011; Tcw et al., 2017). We, therefore, utilized BMP4, and a combination of leukemia-inhibitory factor (LIF), and ciliary-derived neurotrophic factor (CNTF) to activate JAK-STAT signaling, enhancing gliogenesis.

This combination of small molecules, loaded at specific times (**Fig. 1A**), allows efficient conversion of human adult fibroblasts (72-year-old female) to mature astrocytes (**Fig. 1B**). Glial induction was characterized by a change in cellular morphology from typical fibroblast-shaped cells to cells with elongated, bipolar shapes with thinner processes (**Fig. S1**), as well as a concurrent increase in cell number that allowed passaging of cells around D6 of conversion. We hypothesize that the primary cause of this amplification was the initial administration of 2% serum, which, similar to iPSC-iAs, seems to allow a preferential proliferation of astrocytic cells. By around D12, cells reach a distinct phase-dense morphology, likely indicative of a pool of early glial progeny (**Fig. S1**), prompting us to change to media promoting astrocyte differentiation. Critically, albeit limiting cell proliferation, the serum depletion in the remaining stages of the protocol doesn’t seem to increase neurogenesis (**Fig. S2,** MAP2 negative population), indicating that the initial wave of serum is sufficient to alter cell fate towards glia. Additionally, repeated cell-passaging is thought to favor gliogenesis over neurogenesis (Alisch et al., 2021), suggesting that the combination of the two likely assisted in promoting glial conversion. Since GSK-3β inhibition is known to promote neuronal conversion in iPSC studies (Ladewig et al., 2012), we eliminated CHIR99021 during differentiation and maturation of astrocytes. Further, in congruence with Tian et. al., who showed better conversion efficiency with TGF-β inhibitor SB-431542 over A-83-01 (Tian et al., 2016), we eliminated the use of the latter during maturation. Finally, to allow further differentiation, we increased concentrations of LIF and CNTF, which seemed to promote a more star-like morphology, with multiple fibrous branches typical of astrocytes, by about D20 (**Fig. S1**).

**Figure 1.**
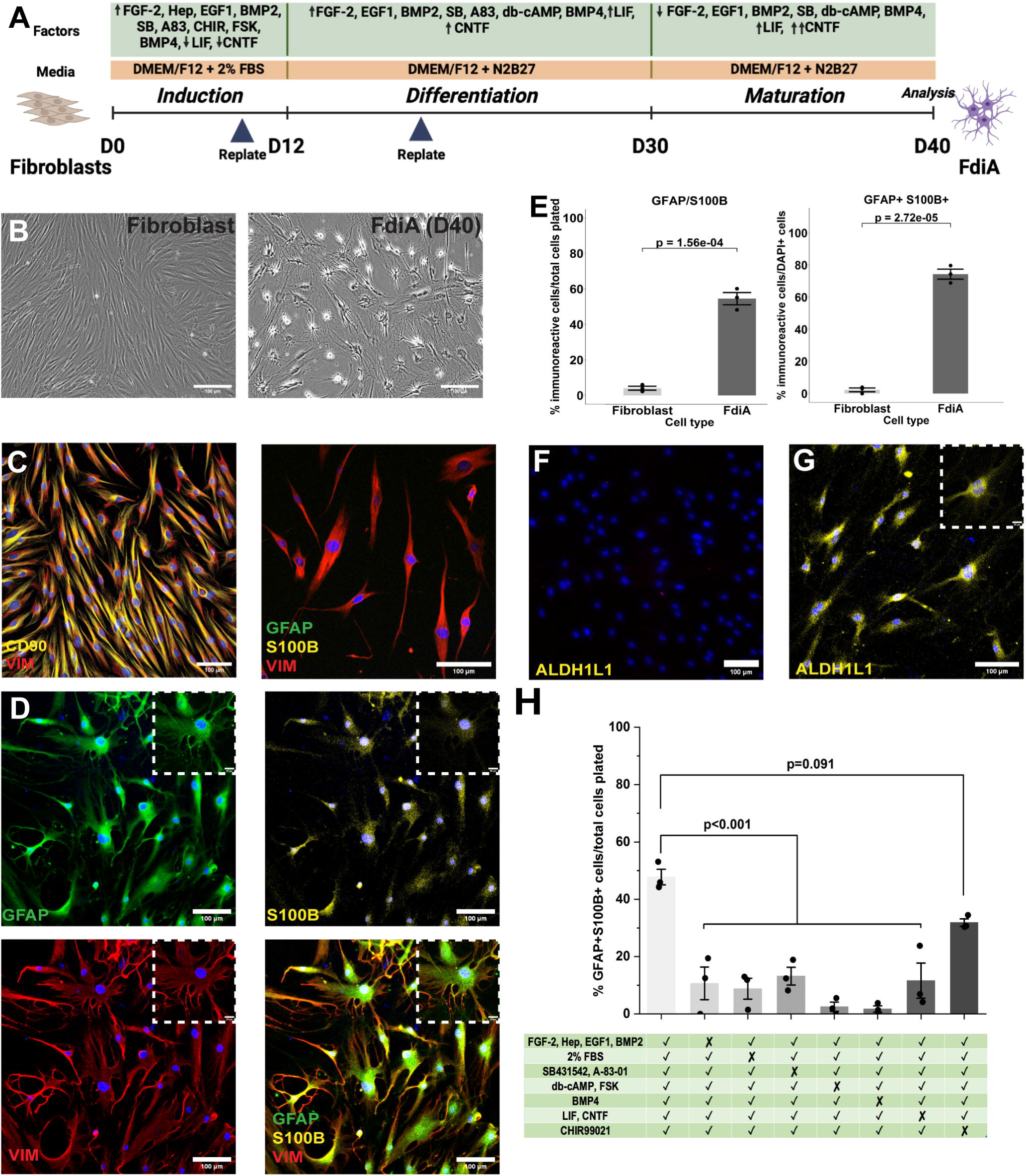
Human adult fibroblasts can be directly converted to induced astrocytes using a combination of small molecules. (A) Experimental overview of the direct conversion protocol to generate Fibroblast-derived induced Astrocytes (FdiAs). *Generated using Biorender.* (B) Phase contrast images of fibroblasts at D0 (left) and FdiAs at D40 (right). (C) Representative immunofluorescence image of CD90/VIM staining (left), and GFAP, S100β and VIM staining (right) in parent fibroblasts at D0. (D) Representative confocal images of GFAP, S100β, VIM in FdiAs at D40 of differentiation. Insets show respective higher magnification images at 63x. (E) Quantification of GFAP, S100β, and GFAP/S100β immunoreactive cells at D40 of differentiation related to the total number of cells (left) and relative to number of DAPI+ cells (right). Data represent Mean ± SEM from n=3 independent experiments, >500 cells. Each dot represents data from individual replicates. Two-tailed independent t-tests were performed for statistical analysis. (F-G) Representative confocal image of ALDH1L1 staining in Fibroblasts (D0). and FdiAs (D40). Insets show respective higher magnification images at 63x. (H) Quantification of GFAP/S100β immunoreactive cells at D40 of differentiation in media deprived of specific small molecules. Data represent Mean ± SEM from n=3 independent experiments, 100-200 cells. Each dot represents data from individual replicates. One-way ANOVA with Tukey’s post-hoc test was performed for statistical analysis (**, p<0.01, ***; p<0.001). All scale bars are 100µm.

Since *SOX9* activation is predominantly required for the initiation of gliogenesis (Kang et al., 2012), we further decreased FGF-2 concentrations and boosted CNTF at day 30 to allow better maturation of the FdiAs. Beyond this stage, astrocytes showed better development of processes, and enlargement of cell bodies until they reach their final maturation state by around D40 (**Fig. S1**). As expected, fibroblasts stained positive for CD90 and Vimentin but were negative for bona fide astrocyte markers, GFAP and S100β (**Fig. 1C**). Our conversion yielded GFAP, S100β, and VIM-positive astrocytes at D40 (**Fig. 1D, 1E**), with ~80% of cells exhibiting dual GFAP/S100β staining per field of view. Taking cell loss during differentiation into account, we estimated ~45% efficiency of conversion based on dual GFAP/S100β staining compared to initial plated cell count (**Fig. 1E**), which is much higher than that observed in any previously published protocols for adult human fibroblasts. FdiAs at this stage, but not fibroblasts, stained for ALDH1L1, a marker for mature astrocytes (**Fig. 1F-G**). We also did not observe any staining of FdiAs for fibroblast (CD90), pluripotency (NANOG, OCT4), neural progenitor (SOX2, PAX6), or neuronal (MAP2) markers (**Fig. S2)**.

Finally, to test whether each pathway proposed above was necessary for conversion, we eliminated small molecules relevant to individual pathways and carried out the protocol as before (**Fig. 1H, S3)**. Removal of any molecule decreased conversion efficiency to <20%. Specifically, the removal of SOX9 inducers appeared to result in a decrease in cell proliferation which could, in part, be attributed to the loss of growth factors (i.e., FGF2 and EGF1). Ablation of SOX9-inducing compounds also resulted in cells lifting from the plate, suggesting cell survival beyond D20 of the conversion protocol was severely impacted. Also, the removal of serum during induction resulted in reduced proliferation, eliminating the need for passaging in the protocol, subsequently decreasing the number of GFAP/S100β positive cells. The elimination of TGF-β inhibitors resulted in the formation of a more rounded morphology, distinct from that observed in the original protocol, staining predominantly to only GFAP and not S100β, suggestive of a potential radial-glia-like intermediate. Removal of cAMP inducers, FSK and db-cAMP, on the other hand, prevented fibroblasts from changing from their initial morphology into glial-like cells. This supports studies from Tian et al. that indicate the absence of OAC1, a small molecule with similar targets as cAMP, limits the conversion ability from one cell type to another (Tian et al., 2016). As expected, removal of either BMP4 or JAK-STAT signaling molecules hindered astrocytic cells from reaching a fully mature state; cells at D40 showed either minimal or no S100β staining while expressing GFAP. Elimination of GSK-3β inhibitor, however, didn’t seem to drastically alter conversion efficiency and was contradictory to previous observations (Tian et al., 2016). We predict that while GSK-3β inhibition may be necessary, it may be limited to a very small window and is unlikely necessary for the entire conversion. Additionally, it is possible that GSK-3β inhibitors only play a role in enhancing the effect of other pathways involved in astrogliogenesis and are, at best, indirect players in the conversion process. Nevertheless, since we still observed a decrease in efficiency, we continued using CHIR99021 during induction for the remainder of this study.

### Human Adult FdiAs express astrocyte-specific transcriptional profiles

To verify that FdiAs are a distinct cell population and resemble astrocytes, we performed bulk RNA-sequencing of the parent fibroblasts and their corresponding FdiAs at D40. Differential expression analysis shows upregulation of several glial lineage genes in FdiAs, including *GJA1, NFIA, NFIB, APOE, SLC1A3* and *SLC16A4,* and downregulation of fibroblast-related genes such as *VIM*, *TGFB1I1* and *COL1A1* (**Fig. 2A, Table S1**). Traditional astrocyte markers, *GFAP* and *S100*β showed either statistical significance (*GFAP*) or sufficient fold change (*S100*β) in FdiAs compared to fibroblasts but did not meet our overall cut-off criteria (log2FC > 1.5, and adj. p-value<0.05) and were excluded from the final set of differentially expressed genes. Similarly, mature astrocyte genes, *AQP4* and *ALDH1L1,* had low read counts across all samples and were filtered in the initial stages of the gene expression pipeline. Larger sample sizes and/or more specific gene isoform analysis might be necessary to validate our immunostaining results, which demonstrated considerable protein expression in FdiAs compared to fibroblasts (**Fig. 1D, 1F, 1G**). To supplement this and verify if our final pool consisted of a mixed cell population, we performed single-cell RNA-sequencing using a combinatorial indexing approach (sc-RNA-seq) and assessed gene expression signatures at different time points during the conversion (**Fig. 2B, S4A**). UMAP plots show cells to predominantly group into four clusters: Fibroblasts (top left cluster) start to rapidly lose their cell-specific expression profile and enter an intermediate state (bottom left), following which they continue to undergo gliogenesis, and form a separate cluster by D10 (bottom right). This is also evidenced by increased expression of early astrocyte marker, *CD49f*, encoded by *ITGA6* (**Fig. S4B**) in the D10 cluster, previously known to be expressed in human iPSC-iAs and fetal brain (Barbar et al., 2020).

**Figure 2.**
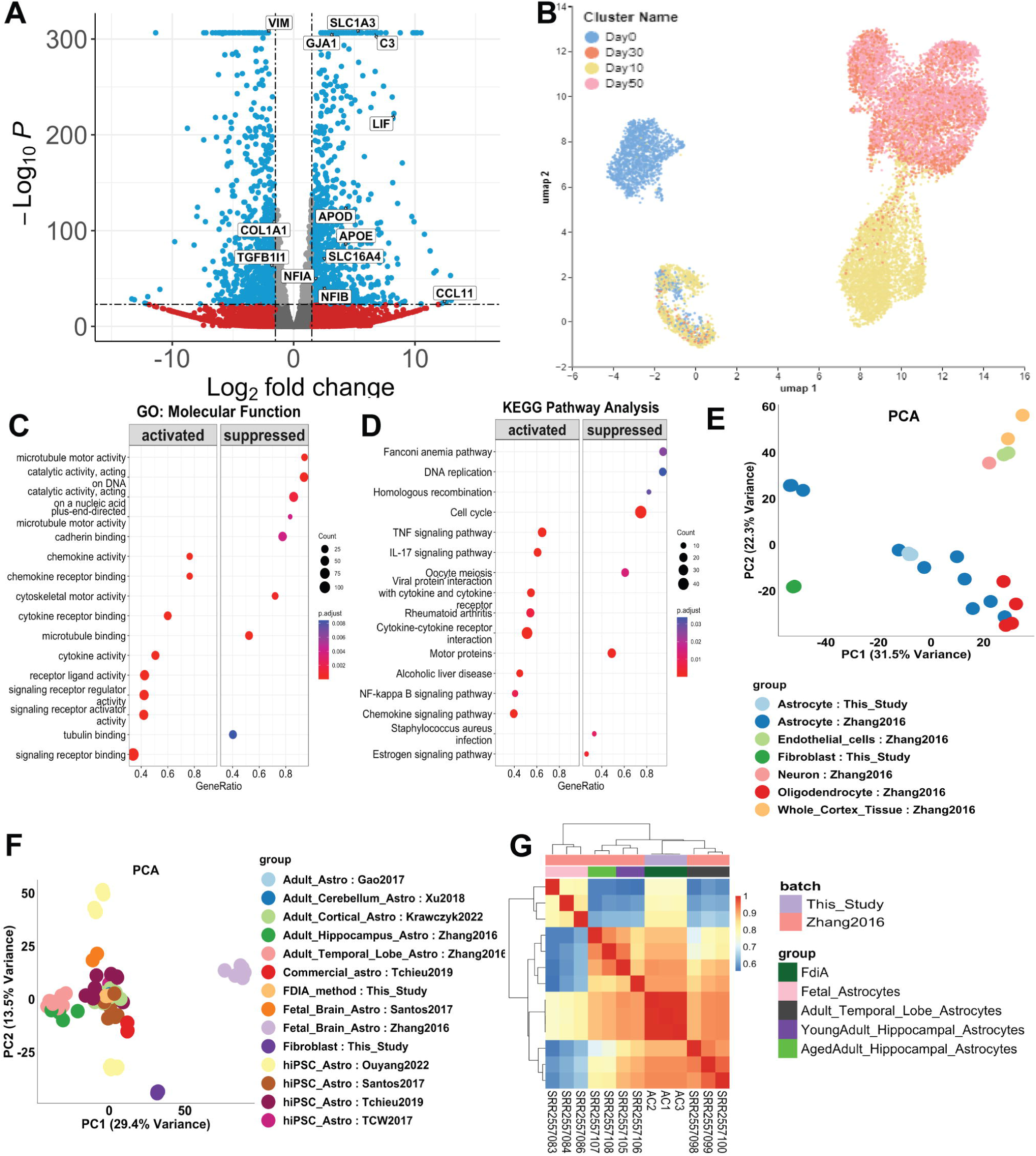
Human adult FdiAs are transcriptionally similar to adult astrocytes. (A) Volcano plot of differentially expressed genes between fibroblasts (D0) and FdiAs (D40), log2FC >|1.5|. (B) UMAP clustering analyses of sc-RNAseq data obtained from cells at D0 (Fibroblasts), D10, D30, and Day 50 of conversion. (C-D) Gene Ontology (enriched for top molecular functions) and KEGG pathway analyses of differentially expressed genes in FdiAs enriched for top molecular functions. (E) Principle component analysis of top 1000 variable genes in fibroblasts and FdiAs (this study), and multiple brain cell types(Zhang et al., 2016). (F) Principle component analysis of top 1000 variable genes in fibroblasts and FdiAs (this study), and other published astrocyte datasets. (G) Sample distance heatmap of all astroglial gene expression data from this study, and Zhang et. al. 2016, clustered based on hclust.

Beyond D10, cells continue to undergo transcriptomic changes and transition into a final cell state, represented by overlapping cell clusters of D30 and D50 FdiAs (top right) (**Fig. S4A**). We next investigated gene expression signatures between this final cluster (D30/50 FdiA), and the starting fibroblasts (D0). Although, consistent with our bulk RNA-seq data, common astrocyte markers including *GFAP* and *S100B* failed to show significantly high expression in the final FdiA cluster (data not shown), we observed continued overexpression of gliogenic markers, including *NFIA and NFIB* (**Fig. S4C-D**) starting during the transition from D0 to D10, with highest expression by D30. Additionally, the common astrocytic glutamate transporter, *GLAST* (encoded by *SLC1A3*, FC >2), as well as, *SPON1* (FC > 3) showed overall high expression at D30 and D50, as compared to starting fibroblasts, as well as D10 FdiAs (**Fig. S4E-F**). *SPON1*, encoding a secretory protein involved in axonal growth, has been previously identified to be conserved in astrocytes (Yang and Jackson, 2019). Moreover, regional sub-type specific analyses of astrocytes in the adult mouse brain identify *SPON1, ETV5,* and other genes to be enriched in a GFAP-low mature astrocyte cluster (AST3) (Batiuk et al., 2020) which seems to correlate with the D30-50 FdiA population. Neuronal genes, including *DCX* and *TUBB3* do not show any significant expression during the entire conversion process (**Fig. S4G-H**), whereas *CD44,* shows near uniform high expression in both fibroblasts and FdiAs (**Fig. S4I**).

Gene ontology and pathway analysis from the bulk RNA-seq data showed genes upregulated in FdiAs to be correlated with canonical astrocyte-like functionality, showing enrichment of pathways and functions predominantly associated with immune function, neuroinflammation and receptor-based signaling activity, whereas, genes upregulated in the fibroblast population were enriched for replication-related processes such as cell cycle and cell division, and cytoskeletal activity (**Fig. 2D-E**). Overall, our analyses suggest that the fibroblasts and FdiAs are two distinct cell populations, with FdiAs closely resembling a specific astrocyte-sub type, with a functional semblance to glial cells.

To confirm this, we obtained human bulk RNA-seq data of multiple brain cell types (Zhang et al., 2016), and astrocyte-specific gene expression data (see Key Resources Table) from previously published studies and compared gene expression signatures. Principle-component analysis (PCA) of RNA-seq data of fibroblasts and FdiAs from our protocol, and brain-cell type specific bulk RNA-seq data from Zhang et. al., showed that our fibroblasts grouped separately from all other cell types (**Fig. 2E**). D40 FdiAs clustered closest to astrocytes from the bulk-RNA seq data, and were separated from other cell types, including neurons, oligodendrocytes, and endothelial cells. This is in line with our immunostaining (**Fig. S2**) and gene expression data (data not shown) that showed little to no detection of common microglial (*IBA1, CXCR1, CD45)*, oligodendrocytic (*OLIG1, OLIG2, SOX10, NG2*), neuronal (*PAX6, SOX1, TBR2, DCX, NEUROD1, MAP2*) and pluripotency-related (*NANOG, OCT4, SOX2, SSEA1*) markers in the FdiAs. PCA of gene expression signatures of FdiAs with multiple published datasets showed that D40-FdiAs closely resembled primary adult cortical astrocytes and those generated from iPSCs (**Fig. 2F**), strengthening our initial observation that FdiAs are a glial cell type. The study by Zhang and others included both fetal and adult human astrocytes. To identify which sub-population of astrocytes our FdiAs resembled, we performed hierarchical clustering analysis on astrocyte data from the two studies (**Fig. 2G**). Sample distance heatmap analysis clustered our FdiAs closer to adult brain-specific astrocytes (r^2^~0.9) than fetal-derived astrocytes (r^2^~0.75). Also, FdiAs were slightly more correlated to temporal lobe neocortical astrocytes (r^2^~0.92) as compared to hippocampal astrocytes (r^2^~0.89).

Since, in culture, we did not observe any significant proliferation of astrocytes during differentiation and maturation, we investigated levels of KI67, in our D40 FdiAs. Strikingly, FdiAs show no expression of KI67 either in immunostaining experiments (**Fig. S5A-C**) or in RNA-seq (**Fig. S5D**), when compared to D0-fibroblasts. We also observed no increase in cell number post the second replating (**Fig. S5E**), suggesting that FdiAs potentially reach a stage of replicative senescence similar to neurons, and might be more representative of astrocytes present in the adult brain (Colodner et al., 2005).

### Human-adult FdiAs exhibit astrocyte-specific functional phenotypes

Astrocytes maintain a homeostatic environment in the brain. They prevent glutamate excitotoxicity in neuronal cells by taking up excess glutamate in their milieu, thus maintaining efficient synaptic transmission in neurons (Mahmoud et al., 2019). Glutamate uptake is mediated by glutamate transporter proteins such as EAAT1 and EAAT2. D40-FdiAs preferentially stain for EAAT2 and show significantly higher expression compared to parent fibroblasts (**Fig. 3A-B**). To quantitatively test uptake capacity, we loaded fibroblasts and D40-FdiAs with extracellular glutamate for 4-6 hours and evaluated the concentration of glutamate left in the media. Supplementing fibroblasts and FdiAs with extracellular glutamate resulted in over two-fold higher uptake in FdiAs than in fibroblasts (**Fig. 3C**). Selective inhibition of EAAT2 with WAY-213613 slightly reduces glutamate uptake in FdiAs (**Fig. 3C**), suggesting that EAAT2 functions along with other transport proteins in glutamate uptake in FdiAs.

**Figure 3.**
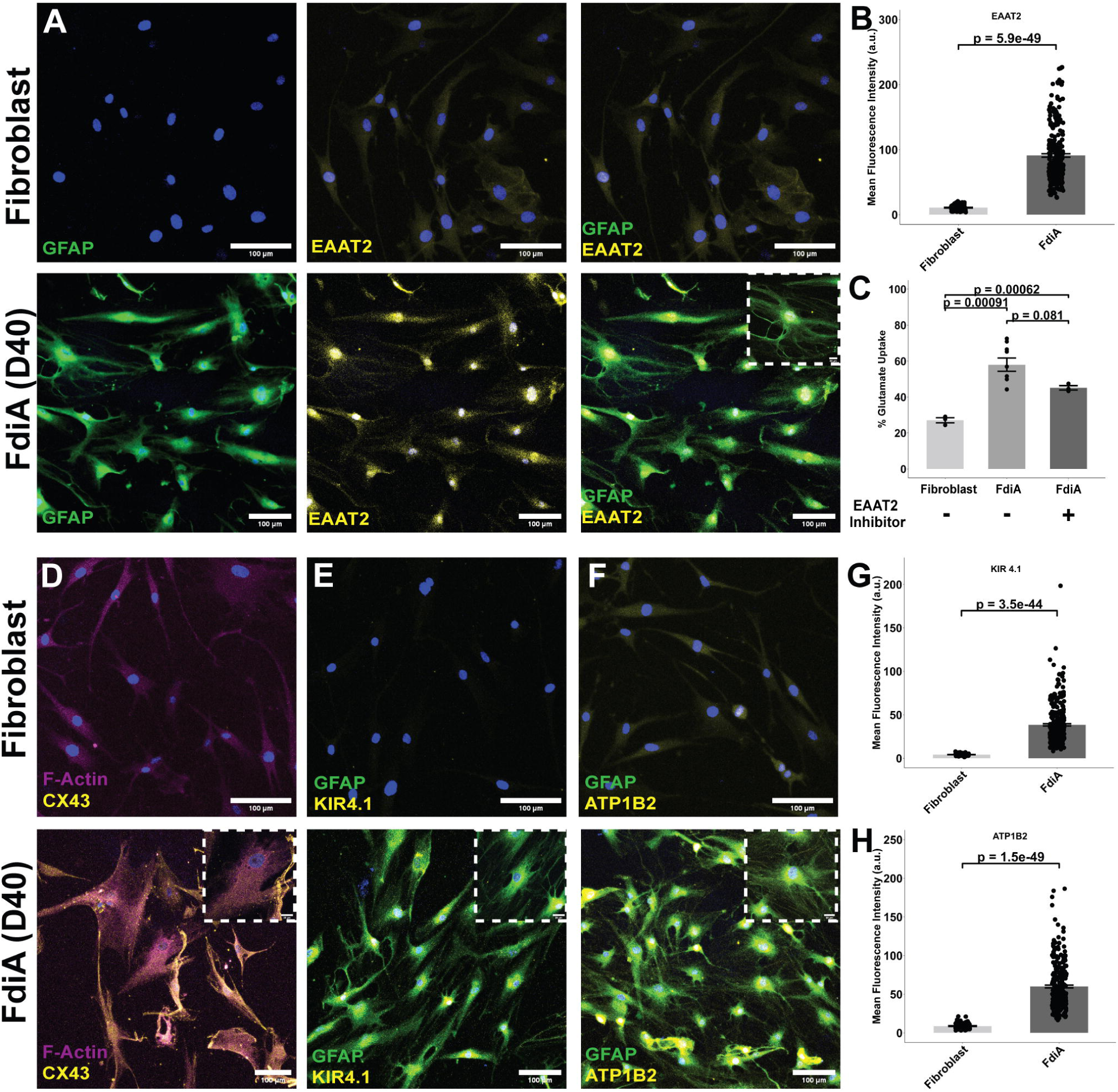
Human adult FdiAs exhibit canonical astrocyte functionality including glutamate uptake, expression of gap-junction proteins, and ionic buffering/homeostasis. (A) Representative confocal images of EAAT2 staining, and its colocalization with GFAP in Fibroblasts (top panel) and FdiAs at D40 (bottom panel). Insets show respective higher magnification images at 63x. (B) Absolute fluorescence intensity measurement of EAAT2 co-stained with GFAP in FdiAs at D40, and fibroblasts (D0). (C) Glutamate uptake capacity of fibroblasts (D0) and corresponding FdiAs (D40). Each dot represents data from individual replicates. (D) Representative confocal images of CX43, co-stained with F-Actin in fibroblasts (top) and in FdiAs at D40 (bottom). Insets show respective higher magnification images at 63x. (E) Representative confocal images of KIR4.1, co-stained with GFAP in fibroblasts and FdiAs at D40. Insets show respective higher magnification images at 63x. (F) Representative confocal images of ATP1B2, co-stained with GFAP in fibroblasts and FdiAs at D40. Insets show respective higher magnification images at 63x. (G-H) Absolute fluorescence intensity measurement of KIR4.1 and ATP1B2 co-stained with GFAP in FdiAs at D40, and fibroblasts (D0). Each dot represents individual cells in all fluorescence intensity analyses (3B,F,H), and data represent Mean ± SEM, for n=150-200 cells, obtained from 10-12 fields of view from three independent replicates. Mann-Whitney U-test was used to assess statistical significance between Fibroblasts and FdiAs in EAAT2, KIR4.1, and ATP1B2 fluorescence intensity measurements. Two-tailed independent t-tests were performed for statistical analysis in all other comparisons. All Scale bars are 100µm.

The ability to form gap junctions and a neuron-glia syncytium allows astrocytes to communicate with other brain cell types, aid in metabolite exchange, and help in rapidly and synchronously responding to changes in CNS stimuli (Liang et al., 2020). Gap junctions are typically composed of proteins called connexins, and astrocytes are known to widely express connexin-43. FdiAs express high levels of CX43 compared to parent fibroblasts (**Fig. 3D, S5A**) and show significantly (log2FC>2) higher gene expression of the CX43 encoding gene, *GJA1* (**Table S1**).

In addition to maintaining homeostatic environments by glutamate regulation and syncytium formation, astrocytes are known to act as potassium ion shuttles, actively contributing to the regulation of neuronal activity (Chever et al., 2010). Specifically, astrocytic expression of KIR 4.1 is known to support the restoration of potassium ion gradients in neurons post-activity (Verkhratsky et al., 2020). Also, astrocytes widely express the Na+/K+ transporter protein, ATP1B2 (Batiuk et al., 2017), known to contribute to similar ionic buffering. To verify whether FdiAs express such proteins, we stained fibroblasts and D40-FdiAs with KIR4.1 and ATP1B2 **(Fig. 3E-H, S5B-C**). FdiAs express nearly 3-4-fold higher levels of KIR4.1, and 2–3-fold higher levels of ATP1B2 compared to unconverted fibroblasts, suggesting that the FdiAs have an in-built machinery to allow this buffering action, like that seen in native human astrocytes. It is likely that expression of these genes indicates functionally mature astrocytes in our cultures, which are capable of maintaining CNS environment-like homeostasis.

Next, we examined whether FdiAs exhibit calcium signaling activity. Previous iPSC-iA studies show that astrocytes exhibit spontaneous and evoked calcium responses (Tcw et al., 2017). We loaded fibroblasts and FdiAs, with Fluo4-AM and recorded calcium response at baseline, and in response to an extracellular pulse of 100µM ATP (**Fig. 4**). FdiAs exhibit a distinct increase in calcium levels upon ATP stimulation compared to parent fibroblasts (**Fig. 4A-C**) and had a significantly higher number of responding cells (~70%) (**Fig. 4D**). Peak amplitudes after ATP stimulation were 3-fold higher than that observed in parent fibroblasts (**Fig. 4E**). This is in line with previous studies of human-embryonic fibroblast-derived astrocytes (Quist et al., 2022) that show significantly higher ATP-evoked responses compared to fibroblasts and primary human fetal astrocytes. Our study extends this to adult human FdiAs, suggesting that intracellular calcium signaling events are successfully captured in direct conversion models. Finally, to validate whether FdiAs can support neuronal cells, we cultured iPSC-derived neurons (**iPSC-iNs)** obtained from the same individual, either alone, or on D40-FdiAs in a co-culture system. We stained neurons for MAP2 and SYN1 (**Fig. 4F-G**). Quantification of the number of MAP2+ neurites, and the number of SYN1+ puncta shows that iPSC-iNs in co-culture with FdiAs express a significantly higher number of SYN1+ puncta (**Fig. 4H**) and show significantly longer neurites (~2-fold) than iPSC-iNs alone (**Fig. 4I**), supporting the idea that FdiAs, like astrocytes *in vivo*, provide a neuronal support environment.

**Figure 4.**
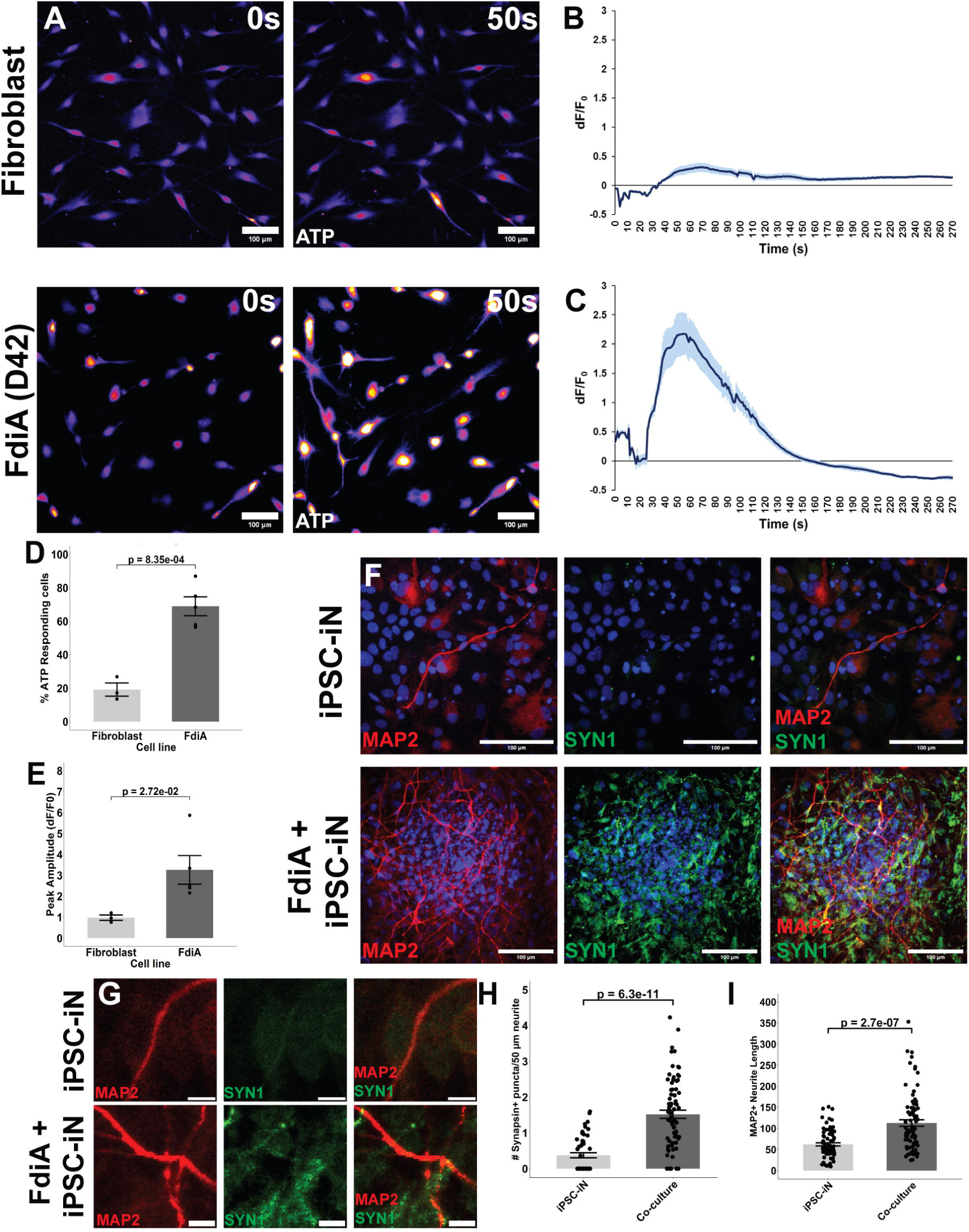
Human adult FdiAs show spontaneous and evoked calcium responses, and support neuronal maturation. (A) Representative background-subtracted images of live-cell calcium imaging experiments with Fluo-4 AM loaded fibroblasts (D0; top panel) and FdiAs (D42; bottom panel) at baseline (0s, left panel) and after ATP stimulation (50s, right panel). All Scale bars are 100µm. (B-C) Mean relative fluorescence intensity changes of calcium traces during ATP-induction experiments in fibroblasts (D0) (B) and D42-FdiAs (C) normalized to baseline (dF/F0). Traces represent Mean ± SEM. (D) Quantification of the percentage of cells responding to ATP stimulus in fibroblasts and FdiAs. Each dot represents data from individual replicates. Two-tailed independent t-tests were performed for statistical analysis. (E) Quantification of the Mean peak amplitude of calcium fluorescence intensity upon ATP stimulation in fibroblasts and FdiAs. Each dot represents data from individual replicates. For calcium imaging experiments, data represent Mean ± SEM, for n=5 independent experiments. Two-tailed independent t-tests were performed for statistical analysis. (F) Representative confocal images of MAP2, co-stained with SYN1 in iPSC-iNs alone (top) and in co-culture with FdiAs (bottom). Scale bars are 100µm. (G) Representative zoomed-in confocal images of MAP2, co-stained with SYN1 in iPSC-iNs alone (top) and in co-culture with FdiAs (bottom). Scale bars are 10µm. (H) Quantification of the number of Syn1 positive puncta per 50µm Neurite in iPSC-iNs and in the FdiA + iPSC-iN co-culture. Each dot represents data from individual neurites. Data represents Mean ± SEM, for n = 50-75 neurites, obtained from 10-12 fields of view from three independent replicates. Mann-Whitney U-test was used to assess statistical significance. (I) Quantification of the length of MAP2+ neurites in iPSC-iNs and FdiA + iPSC-iN co-culture. Each dot represents data from individual neurites. Data represents Mean ± SEM, for n = 50-80 neurites, obtained from 10-12 fields of view from three independent replicates. Mann-Whitney U-test was used to assess statistical significance.

### FdiAs are immunocompetent and respond to cytokine stimuli

Astrocytes have been long known to respond to microglial cytokine stimuli, an effect that sees their conversion from an ischemic A2 state to a pro-inflammatory A1 reactive state (Giovannoni and Quintana, 2020). Similar properties have been previously observed in both iPSC-derived and fibroblast-derived astrocyte models (Liddelow et al., 2017; Quist et al., 2022; Tcw et al., 2017). To mimic microglial neuroinflammatory stimuli, we loaded D40-FdiAs with a cocktail of three cytokines (TNFα, IL-1α, and C1q) for 24 hours and performed bulk RNA-seq on stimulated cells. Differential gene expression analysis between stimulated FdiAs and unstimulated controls at D40, unsurprisingly, showed a significant upregulation in genes related to a neuroinflammatory phenotype (**Fig. 5A**), including responses to cytokine stimulation and IFN-γ activation, typical of astrocytes, as was evident from gene ontology and pathway analysis (**Fig. 5B-D**). We also observed a concomitant downregulation of genes related to homeostasis maintenance properties of astrocytes, such as potassium ion regulation, pointing to a loss of ischemic phenotype in the stimulated FdiAs. Cytokine-stimulated FdiAs, in fact, exhibited >3-fold increase in several cytokines, known to be indicative of the A2 phenotype, including *IL-1A, IL-1B, CXCL2, CCL20, C3,* and others, thereby suggesting that our model can potentially switch between reactive states, and could be used to model neuroinflammation in age-related brain disorders.

**Figure 5.**
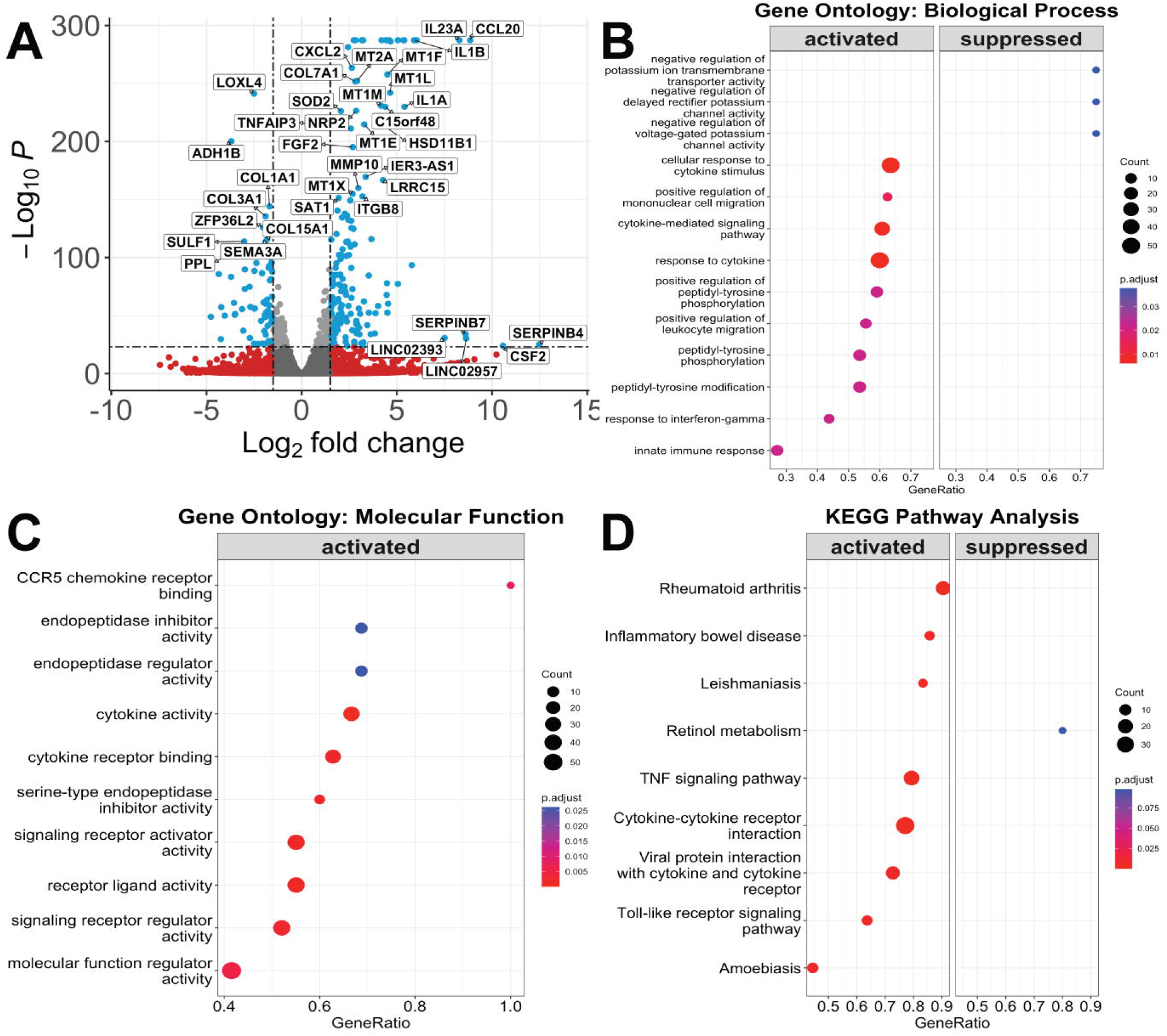
Human adult FdiAs are immunocompetent. (A) Volcano plot of differentially expressed genes between control FdiAs at D40 and cytokine-stimulated FdiAs at D40, log2FC >|1.5|. (B) Gene Ontology analyses of differentially expressed genes in stimulated-FdiAs enriched for top biological processes. (C) Gene Ontology analyses of differentially expressed genes in stimulated FdiAs enriched for top molecular functions. Note: No suppressed molecular functions showed statistical significance in this data. (D) KEGG pathway analysis of enriched pathways amongst differentially expressed genes in cytokine-stimulated FdiAs.

### Direct conversion yields FdiAs irrespective of age and sex of starting fibroblasts

For any *in vitro* model to be applicable at large, factors such as age and sex must be considered. Previous studies on FdiAs indicated the ability of fetal fibroblasts to be converted to astrocytes with high efficiency (Quist et al., 2022; Tian et al., 2016), and our experiments expanded this conversion capability to adult fibroblasts derived from a single individual. We, therefore, tested whether this conversion strategy could be effectively applied to other fibroblast populations by obtaining fibroblasts from two additional healthy individuals: a young female (20 years old) and a healthy aged male (72 years old). We converted these fibroblasts to FdiAs using our standardized protocol (**Fig. 1A**) and observed that by D40, both cell lines exhibited significantly higher GFAP and S100β expression than their corresponding parent fibroblasts (**Fig. 6A**). Importantly, conversion efficiency was highly comparable to our previous cell line, ranging between 38-50% (**Fig. 6B**), supporting the efficacy of our method. Additionally, we observed a remarkably similar ability of these FdiAs to take up glutamate, as observed in our other cell line (**Fig. 6C**).

**Figure 6.**
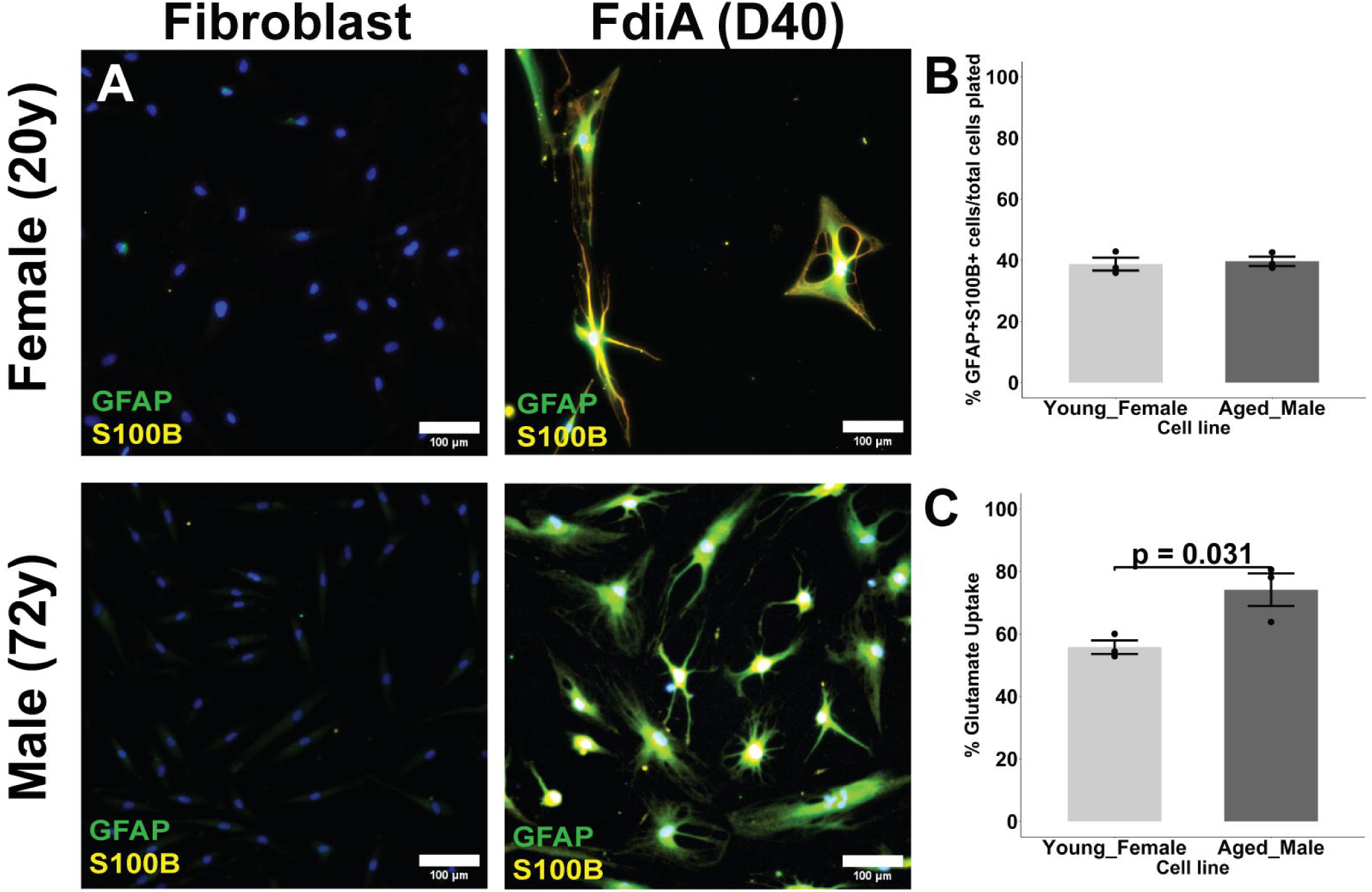
Generation of human adult FdiAs from a healthy young female fibroblast and healthy aged male fibroblast. (A) Representative immunofluorescence images of GFAP and S100β staining in Fibroblasts (D0) and FdiA (D40) from one healthy young female (top panel), and one healthy aged male (bottom panel). (B) Quantification of GFAP and S100β immunoreactive cells at D40 of differentiation normalized to the total number of cells. (C) Glutamate uptake capacity of the two FdiAs (D40). Each dot represents data from individual replicates. All data represent Mean ± SEM from n=3 independent experiments, >150 cells. Two-tailed independent t-tests were performed for statistical analysis. All Scale bars are 100µm.

Taken together, these experiments suggest that our improved method reliably and consistently generates functional, mature astrocytes from human adult fibroblasts at high efficiency and that it is broadly applicable to individuals of different age and sex.

## Discussion

Stem-cell-derived models have long been the forerunners in the study of diseases in a dish. However, in recent years, several studies point to an inherent inability of iPSC-derived models to capture age-associated changes and genetic mosaicism of parent cells, and their resemblance to embryonic or pre-natal cell types (Mertens et al., 2015). Also, the generation of mature cell types like astrocytes from iPSCs is known to be a time-intensive process. As an alternative, and to supplement studies of iPSC-derived models, multiple groups have generated neurons, oligodendrocytes, and other cell types directly from a starting population of fibroblasts that seem to retain adult-specific epigenetic and transcriptomic markers (Koch, 2015; Li et al., 2015; Mertens et al., 2015). However, similar approaches have been lacking for astrocytes, and to our knowledge, only three prior studies have established the generation of iAs directly from fibroblasts (Caiazzo et al., 2015; Quist et al., 2022; Tian et al., 2016). While all three of these studies were able to obtain iAs at high efficiency from either mouse or human fibroblasts of fetal origin, they were unable to maintain similar conversion efficiencies for human fibroblasts of adult origin. Although studies of direct conversion from fetal fibroblasts are relevant, they likely only capture pre-natal developmental stages and are unlikely to be suitable for modeling sporadic late-onset neurological disorders.

In this study, we developed a highly improved small-molecule-based conversion strategy that can generate iAs from human adult fibroblasts with great efficiency (~40%). We found that this protocol is consistent across age and sex, and that conversion efficiency is significantly higher than that observed in previous studies. Tian et.al., used a cocktail of five small molecules across the entire differentiation process to generate iAs. While three of these compounds also played a crucial role in our conversion method, our use of additional compounds in a timed manner likely activated other pathways that enhanced the process. For instance, Tian et. al. observed maximal conversion efficiency of mouse embryonic fibroblasts to iAs (~38% GFAP+ cells) when using a combination of CHIR99021, SB-431542 (both used in this study), a histone deacetylase inhibitor (VPA), histone demethylase LSD1 inhibitor (tranylcypromine) and *Oct4* activator (OAC1). While the use of GSK3b inhibitor (CHIR99021) and TGF-b inhibitor (SB431542, A-83-01) seems to play a critical role in glial reprogramming in both studies, VPA and OAC1 seemed to only reduce conversion efficiency minimally in the former study (Tian et al., 2016) and were omitted from our study. VPA and tranylcypromine, both histone modifiers, have been shown to activate embryonic stem cell and pluripotency-related genes including *Oct4*, and have previously been used to generate iPSCs from fibroblasts (Huangfu et al., 2008; Lee et al., 2006; Li et al., 2011). The combined use of these compounds in FdiA conversion will likely result in an intermediate pluripotent state that resets the epigenome thereby erasing adult-cell-relevant properties. We therefore excluded these compounds in our study. Importantly, we also activated canonical astrocyte maturation pathways (BMP4, and JAK-STAT), in line with the approach used by Quist et. al., which was lacking in the study by Tian et. al. Our approach to using small molecules in place of lentiviral expression of transcription factors avoids the unnecessary need to integrate foreign DNA and offers an easy-to-implement method, in a reduced time frame.

In addition to morphological assessments and immunofluorescence assays that show our FdiAs to have astrocytic morphology and express mature astroglial markers (S100β, ALDH1L1, EAAT2), we demonstrated the presence of gap-junction (CX43) and ion-channel (KIR4.1, ATP1B2) proteins that highlight the potential ability of FdiAs to maintain ionic and buffering homeostasis. Our FdiAs also exhibit efficient glutamate uptake ability, calcium signaling activity, provide neuronal support, and show response to cytokine stimuli, bolstering our interpretation that these FdiAs are of an astroglial lineage. Critically, in comparing FdiAs obtained from human *adult* fibroblasts, our protocol results in higher conversion efficiency (~80% GFAP+/S100b+ cells) than previous studies using human adult fibroblasts for iA conversions (8-10% GFAP+ or S100B+ cells, (Tian et al., 2016)). Our FdiAs also exhibit better glutamate uptake (~60%) than human adult FdiAs in the Quist et. al. study (~20-25%), which is the only prior study that assessed this (Quist et al., 2022).

RNA-seq is a powerful method to evaluate relevant cellular signatures. However, limited sample sizes in the current study and/or potential rapid RNA degradation may have contributed to an inadequate read-out of some genes known to be over-expressed in astrocytes. Despite this limitation, corroborating our immunofluorescence data, gene expression analysis of FdiAs offered some critical clues to the validity of our model. sc-RNA-seq data from the conversion process shows fibroblast differentiation to occur by ~10 days, with increased expression of early astroglial markers like CD49f, suggesting a successful transition towards glial fate. Comparison with published data sets indicates that FdiAs cluster closest to human adult astrocytes, particularly temporal lobe astrocytes, than other cell types, and exhibit gene expression profiles similar to other *in vitro* astrocyte models. Further, given the overall high expression of genes like *SPON1* observed in our sc-RNA-seq data, it will be interesting to assess if FdiAs mimic a specific sub-population, providing an important step toward the generation of more cortical astrocyte sub-types. Although our study was not designed to assess retention of age via epigenetic and transcriptional profiles, we expect that like iNs, our conversion strategy will provide an aged-astrocyte model that will be highly useful for the study of age-related brain disorders. That said, further analysis with increased sample sizes using fibroblasts that span a wider age group is warranted to bolster our findings.

Overall, our study offers a proof-of-principle, wherein, the efficient generation of functional, mature, astrocytes, directly from human adult fibroblasts, using a cocktail of small molecules, resembles transcriptomic signatures of adult cortical astrocytes, and offers a substantially improved model for the investigation of adult brain disorders.

## EXPERIMENTAL PROCEDURES

### RESOURCE AVAILABILITY

#### Corresponding author

Melanie A. Carless.

#### Material availability

Reagents used in this study are publicly available, or available from the corresponding author upon reasonable request.

#### Data and code availability

The accession number for the bulk RNA-Seq raw data, and sc-RNA-seq data generated in this paper are GSE239320, and GSE272279 respectively.

### EXPERIMENTAL MODEL AND SUBJECT DETAILS

#### Subjects

Both fibroblast and iPSC cell lines were obtained from Coriell (NIA Aging Cell Culture Repository, Baltimore longitudinal study of aging, and NIGMS Human Genetic Cell Repository). **Table S2** contains cell line details.

#### Generation of Fibroblast-derived induced Astrocytes (FdiAs)

Adult human fibroblasts were regularly cultured in DMEM/F12 with 15% FBS and 0.1% NEAA (all Gibco). For direct conversion experiments, fibroblasts were plated on Matrigel (Corning) coated dishes at 1.25×10^4^cells/cm^2^ and cultured for 24h. Conversion followed a three-step process. Starting at D1, media was changed to Astrocyte-induction media (DMEM/F12 with 2% FBS, 1% B-27, 1% NEAA, 1% GlutaMAX, and 1% Penicillin- streptomycin (all Gibco)), supplemented with a small molecule cocktail: 12ng/ml FGF2 (ThermoFisher), 2ug/ml Heparin (Sigma), 5 ng/ml EGF-1 (Peprotech), 200ng/ml BMP2 (Peprotech), 10µM SB-431542 (Sigma), 1µM A-83-01 (Sigma), 3µM CHIR99021 (LC Labs), 5µM Forskolin (LC Labs), 10ng/ml BMP4 (Peprotech), 10ng/ml LIF (Peprotech) and 5ng/ml CNTF (Peprotech). Half-media changes were given on alternate days until D12. Confluent cells were passaged using Accutase (Thermo Fisher), and re-plated at 1.25×10^4^ cells/cm^2^ on Matrigel-coated dishes. On D13, media was changed to astrocyte-growth media containing DMEM/F12 with 1% B27, 1% N2, 1% NEAA, 1% GlutaMAX and 1% Pen-Strep. Small molecule cocktail in growth media was devoid of Heparin, CHIR99021 and FSK but contained 500 ug/ml db-cAMP (Santa Cruz Biotechnology), and higher dosage of LIF (30 ng/ml) and CNTF (10 ng/ml). All other components were identical to those used in the astrocyte-induction media. Cells were given half-media changes every third day from here on. For immunofluorescence and calcium imaging assays, cells were plated on Matrigel-coated sterile glass coverslips. For further maturation, media was changed to astrocyte-maturation media at D30. All components of the maturation media were identical to growth media except for FGF2 (4 ng/ml), CNTF (20 ng/ml) and A-83-01 (removed). All experiments were carried out between 40-50 DIV and detailed methodologies are outlined in **Supplementary Methods**.

#### Statistical Analysis

Statistics were performed in R, using the methods indicated for each figure or as in the Supplementary Methods. Unpaired independent t-tests or Mann Whitney-U tests were used, based on the normality of data (calculated using the Shapiro-Wilk test, p>0.05). For multiple group comparisons, one-way ANOVA was used.

## ACKNOWLEDGEMENTS

This work was supported by the National Institute of Health grant, 1R21AG085428 (MC), the American Federation for Aging Research – Diana Jacobs/Kalman Scholarship (UB), and Institutional start-up funds from UTSA (MC). We would like to thank the UTSA-NDRB Department, Lindsey Macpherson and Gary Gaufo for access to the confocal microscope; and the UTSA Cell Analysis Core for assistance with FACS. Some figures were created using Biorender.com.

## AUTHOR CONTRIBUTIONS

U.B. and M.C. conceived and designed experiments. U.B., E.S., J.A., and A.P. performed cell culture experiments. U.B. and E.S. performed Confocal Imaging. U.B., E.S., J.A., and A.P. conducted image analyses. U.B., M.K., and E.S. performed RNA-seq and sc-RNA-seq analyses. U.B. wrote the manuscript with input from all authors. M.C. edited the manuscript and provided financial support.

