## Supplementary Material for "An Efficient Direct Conversion Strategy to Generate Functional Astrocytes from Human Adult Fibroblasts"

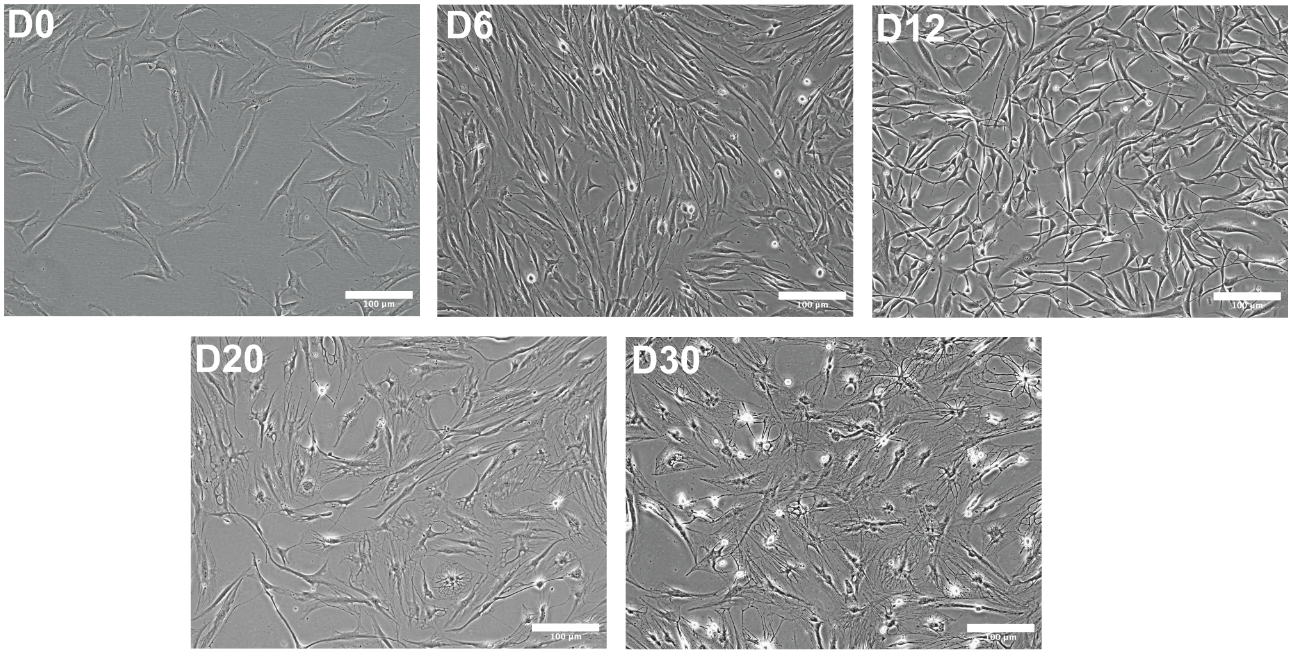
D0
100 µm
D6
100 µm
D12
100 µm
D20
100 µm
D30
100 µm

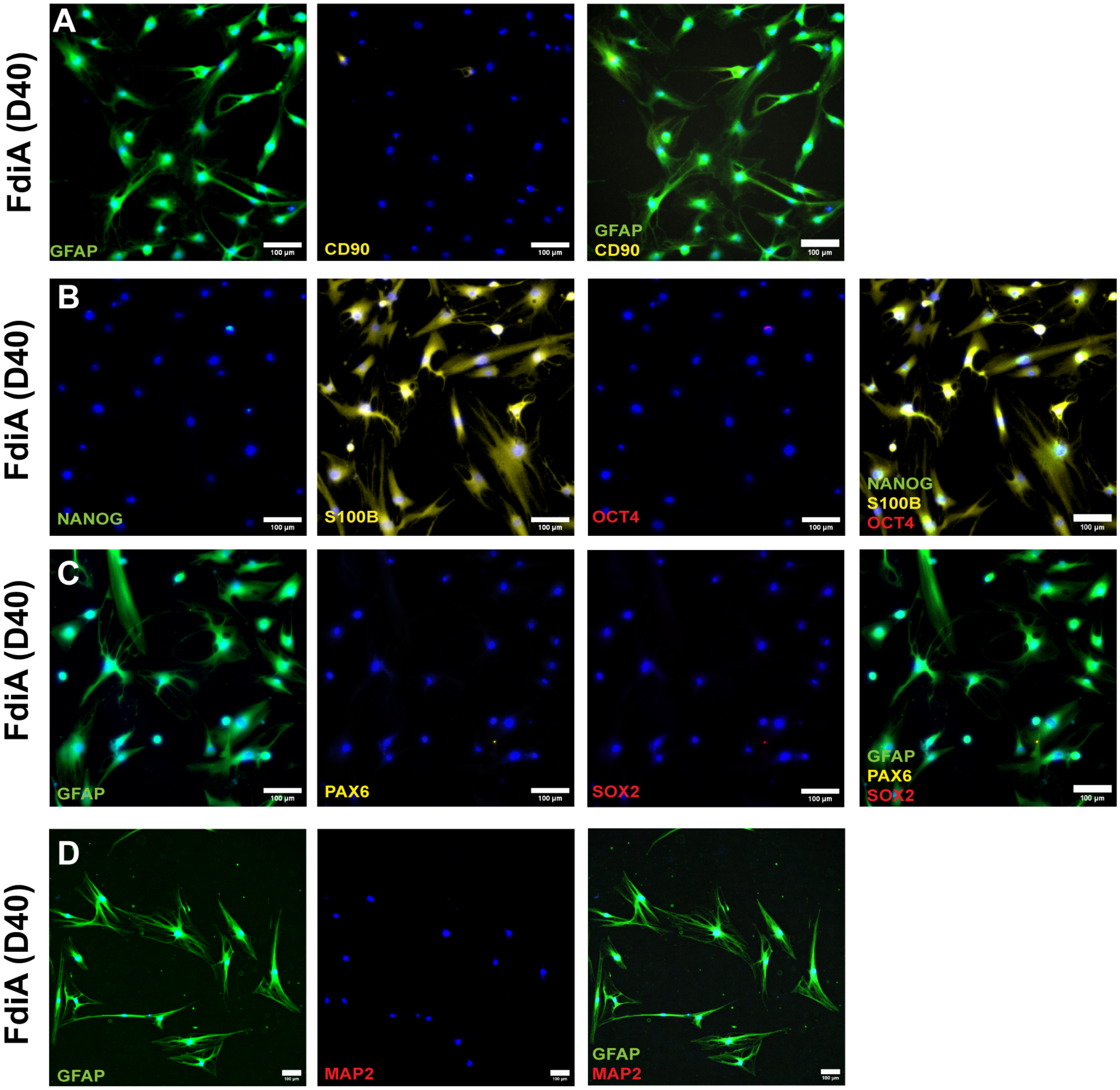

A
FdiA (D40)
GFAP
CD90
GFAP
CD90
B
FdiA (D40)
NANOG
S100B
OCT4
NANOG
S100B
OCT4
C
FdiA (D40)
GFAP
PAX6
SOX2
GFAP
PAX6
SOX2
D
FdiA (D40)
GFAP
MAP2
GFAP
MAP2
100 µm

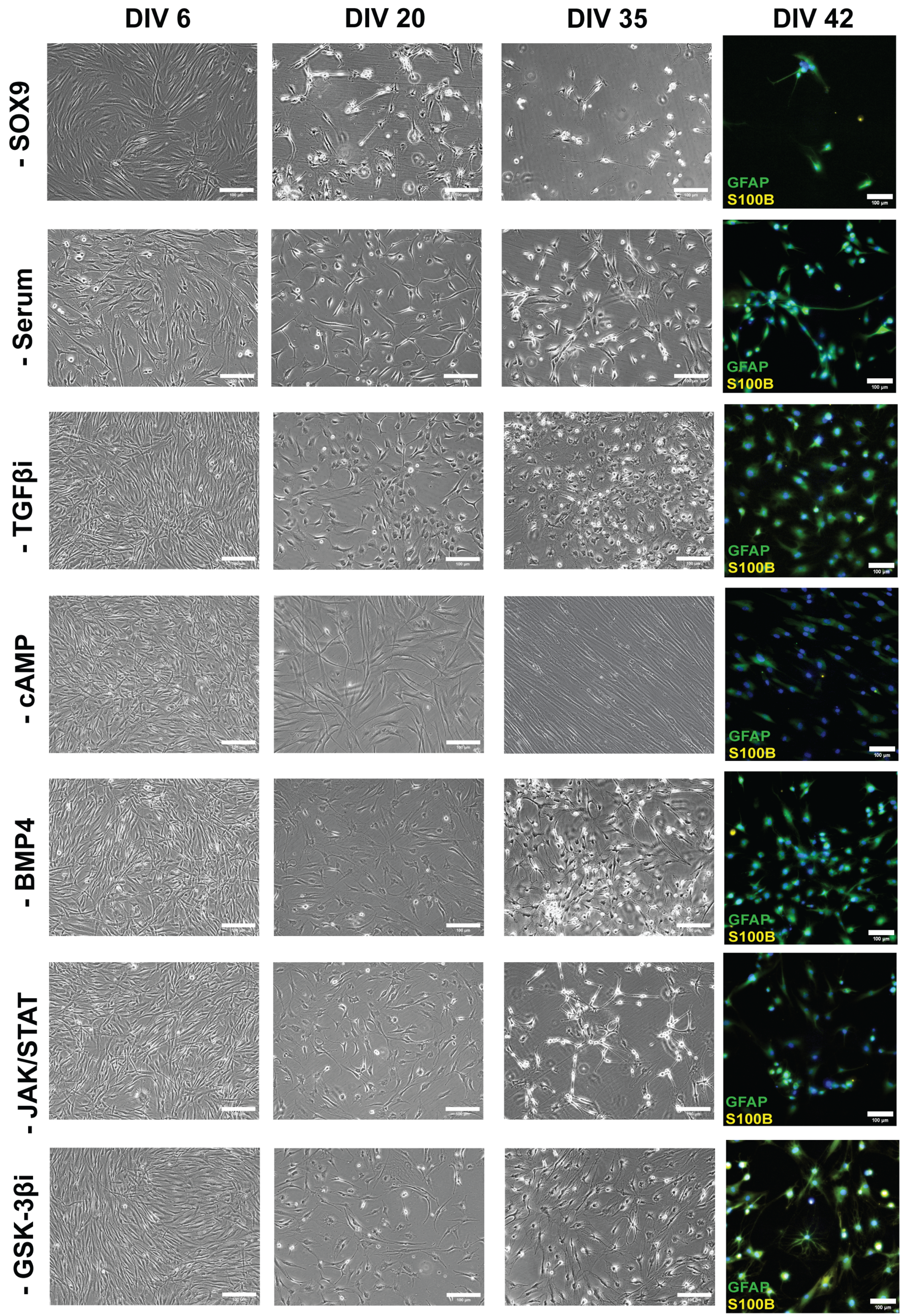
DIV 6
DIV 20
DIV 35
DIV 42
- SOX9
- Serum
- TGFβi
- cAMP
- BMP4
- JAK/STAT
- GSK-3βi
GFAP
S100B

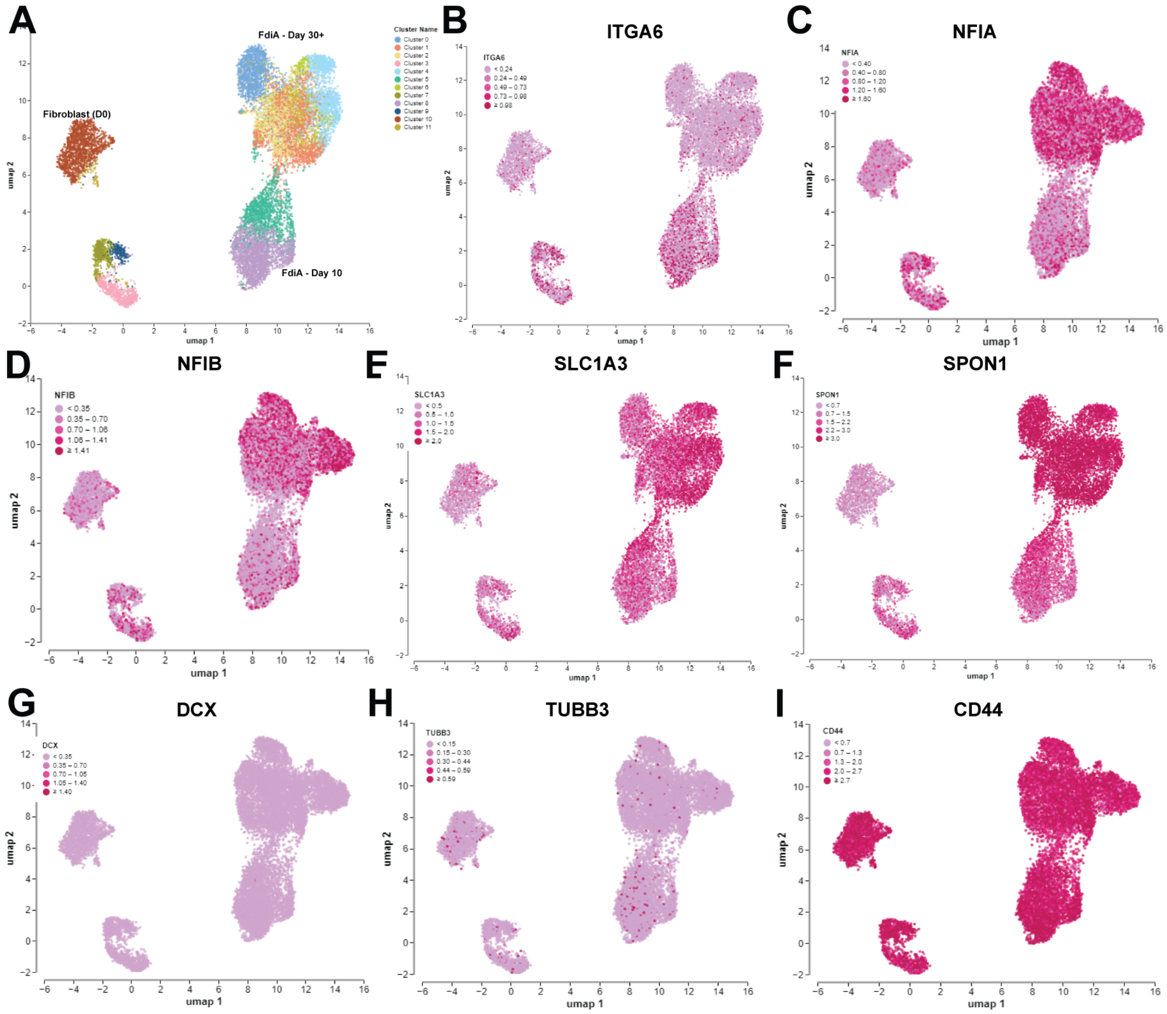
A
FdiA - Day 30+
Fibroblast (D0)
FdiA - Day 10
Cluster Name
B
ITGA6
C
NFIA
D
NFIB
E
SLC1A3
F
SPON1
G
DCX
H
TUBB3
I
CD44
umap 1
umap 2

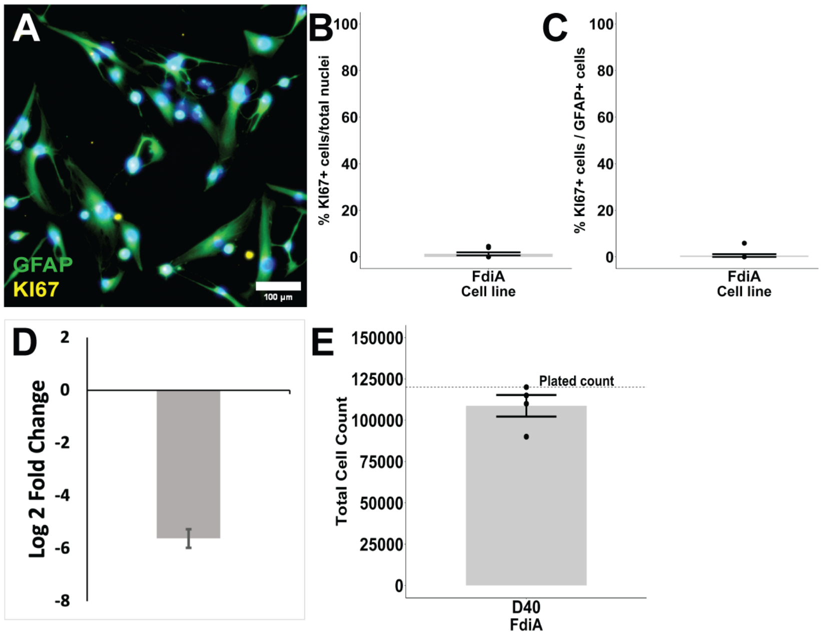
A
GFAP
KI67
100 µm
B
% KI67+ cells/total nuclei
100
80
60
40
20
0
FdiA
Cell line
C
% KI67+ cells / GFAP+ cells
100
80
60
40
20
0
FdiA
Cell line
D
Log 2 Fold Change
2
0
-2
-4
-6
-8
E
Total Cell Count
150000
125000
100000
75000
50000
25000
0
Plated count
D40
FdiA

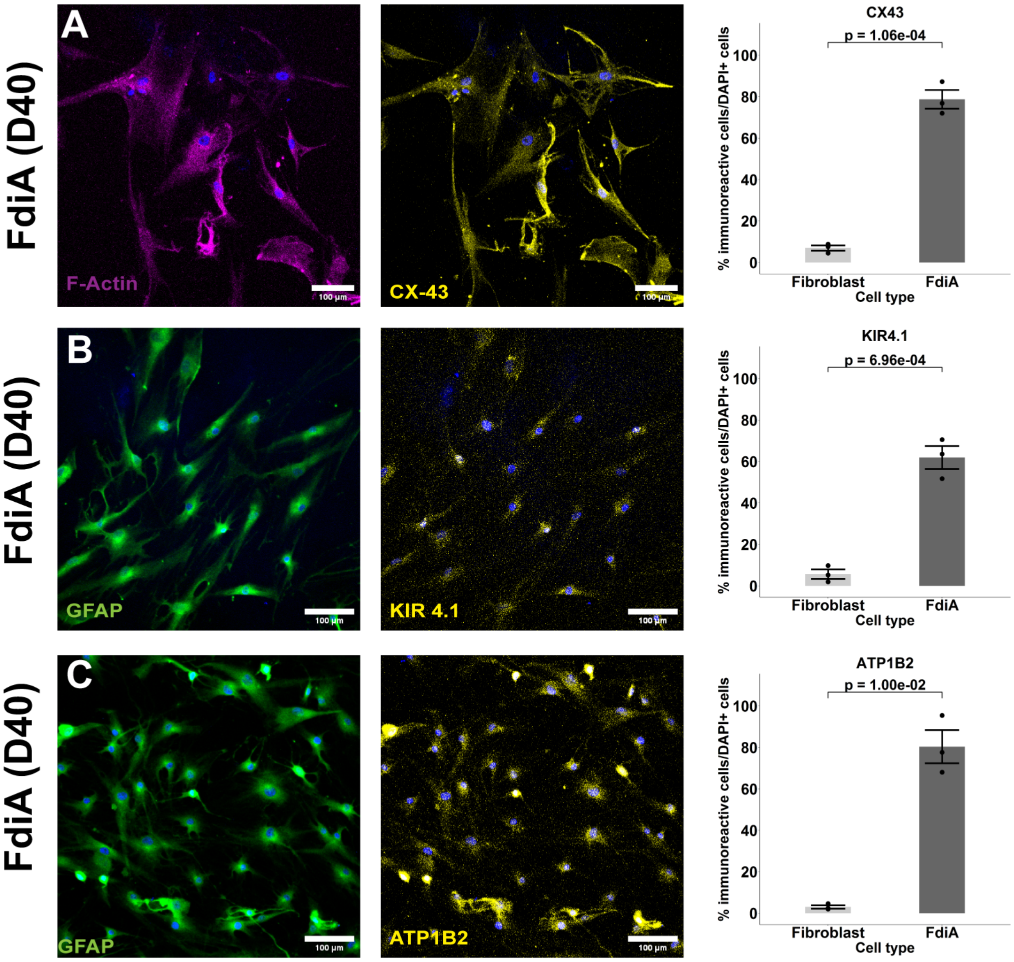

A
B
C
FdiA (D40)
FdiA (D40)
FdiA (D40)
F-Actin
CX-43
GFAP
KIR 4.1
GFAP
ATP1B2
100 µm
CX43
KIR4.1
ATP1B2
p = 1.06e-04
p = 6.96e-04
p = 1.00e-02
% immunoreactive cells/DAPI+ cells
Fibroblast
FdiA
Cell type

#### SUPPLEMENTAL INFORMATION

#### SUPPLEMENTARY FIGURES

**Figure S1. Phase contrast images of the astrocyte-conversion process.** Representative phase contrast images show fibroblasts before conversion (D0), during induction (D6), before switching to differentiation media (D12), during differentiation (D20), and before switching to maturation media (D30). All Scale bars are 100μm.

**Figure S2. Human Adult FdiAs do not express markers of other cell types.** Representative immunofluorescence images of FdiAs stained for fibroblast-specific marker, CD90 (A), pluripotency markers, NANOG and OCT4 (B), neural stem cell markers, PAX6 and SOX2 (C), and neuronal marker, MAP2 (D), co-stained with either GFAP or S100β. All Scale bars are 100μm.

**Figure S3. Establishing a minimal set of small molecules required for direct conversion to FdiAs.** Representative phase contrast (D6, D20, D30) and immunofluorescence images (D42) of GFAP/ S100β dual-stained FdiAs, in *SOX9* depleted, serum depleted, TGF-β activated, cAMP depleted, BMP4 depleted, JAK-STAT pathway inactivated, and GSK-3β activated media. All scale bars are 100μm. (- sign indicates subtraction; and 'i' refers to inhibition. See Fig. 1H for quantification).

**Figure S4. Gene Expression Analysis points to Glial Lineage in FdiAs.**

(A) UMAP plot of sc-RNAseq data obtained during FdiA conversion shows distinct clustering of FdiAs in each stage of the protocol.

(B-I) Representative UMAP clustering plot with log-normalized expression levels from sc-RNAseq data during FdiA conversion for astrocyte-specific genes (ITGA6, NFIA, NFIB, SPON1, SLC1A3), neuronal genes (DCX, TUBB3), and common fibroblast/astrocyte gene (CD44).

**Figure S5. Human adult FdiAs express gap-junction proteins and show ionic buffering/homeostasis.** Representative single-stained confocal immunofluorescence images, and their respective quantification in Fibroblasts and FdiAs for CX43 (A), KIR4.1 (B), and ATP1B2 (C). Data represent Mean ± SEM from n=3 independent experiments, 100-200 cells. Each dot represents data from individual replicates. Two-tailed independent t-tests were performed for statistical analysis. All scale bars are 100μm. *Also see Fig. 3.*

**Figure S6. Human adult FdiAs are not proliferative at D40 of differentiation.**

(A) Representative immunofluorescence images of KI67 and GFAP dual-stained FdiAs at D40.

(B-C) Quantification of the percentage of KI67 immuno-positive cells to the total number of cells (B), and the number of GFAP immunoreactive cells (C), in FdiAs.

(D) Differential expression analysis of *MKI67* gene expression in fibroblasts vs D40 FdiAs. Log2FC represents the downregulation of *MKI67* in FdiAs.

(E) Cell count data of FdiAs after 40 DIV when seeded at an initial density of 120,000 cells per 35mm dish during the second replating. Data represent Mean ± SEM. Mann-Whitney U-test was used to assess statistical significance. All Scale bars are 100µm.

#### SUPPLEMENTARY TABLES

**Table S1.** Differential gene expression of commonly expressed astrocyte and fibroblast markers.

| Marker | Gene | log2 Fold Change* | Padj# |
|---|---|---|---|
| Astrocyte | *LIF* | 8.2675 | 6.49E-221 |
| | *C3* | 6.6840 | 1.00E-06 |
| | *CNTFR* | 6.0985 | 6.18E-05 |
| | *SLC1A3* | 5.4802 | 1.00E-06 |
| | *APOD* | 4.4885 | 1.92E-124 |
| | *APOE* | 4.1114 | 3.33E-83 |
| | *TNC* | 4.0779 | 1.00E-06 |
| | *ATP1B2* | 3.5181 | 2.37E-05 |
| | *GJA1* | 3.3388 | 1.00E-06 |
| | *SLC15A2* | 2.6612 | 5.83E-03 |
| | *S100B* | 2.5686 | n.s |
| | *SLC16A4* | 2.3922 | 5.85E-69 |
| | *NFIB* | 2.3620 | 3.27E-41 |
| | *NFIA* | 1.9749 | 2.04E-48 |
| | *ATF3* | 1.5937 | 2.17E-06 |
| | *SOX9* | 1.5346 | 4.34E-12 |
| | *GFAP* | 1.4320 | 7.47E-04 |
| | *GPC4* | 0.5176 | 7.54E-03 |
| | *BMPR1B* | 0.4067 | 4.61E-01 |
| | *RUNX2* | 0.3524 | 3.03E-02 |
| | *DIO2* | 0.2755 | 6.49E-01 |
| | *ALDOC* | 0.1423 | 6.34E-01 |
| | *F3* | 0.1256 | 3.26E-01 |
| | *SLC25A18* | 0.0162 | 9.89E-01 |
| | *NNAT* | -0.2682 | 8.36E-01 |
| | *NOTCH1* | -1.4901 | 7.81E-28 |
| | *IGFBP7* | -1.6613 | 3.20E-118 |
| | *TUBA1A* | -1.7702 | 1.52E-180 |
| | *ATP1A2* | -3.2153 | n.s. |

| | | | |
|---|---|---|---|
| | *MLC1* | -3.2454 | n.s. |
| | *SLC14A1* | -4.0586 | 1.82E-01 |
| | *FABP7* | N/A | N/A |
| | *KCNJ10* | N/A | N/A |
| | *HEPACAM* | N/A | N/A |
| | *AQP4* | N/A | N/A |
| Fibroblast | *DES* | -4.0526 | 8.97E-03 |
| | *VIM* | -2.2284 | 1.00E-06 |
| | *TGFB1I1* | -1.6000 | 5.01E-61 |
| | *COL1A1* | -1.4104 | 2.59E-110 |
| | *FN1* | -0.7570 | 2.46E-17 |
| | *S100A4* | -0.4739 | 3.18E-07 |
| | *ECM1* | -0.4646 | 1.30E-12 |
| | *FBLN2* | 0.0761 | 4.07E-01 |
| | *ACTA2* | 0.1586 | 1.87E-01 |
| | *COL1A2* | 0.1992 | 4.94E-03 |
| | *PDGFRB* | 0.2549 | 9.03E-03 |
| | *CDH11* | 0.5863 | 1.18E-12 |
| | *POSTN* | 1.2774 | 5.18E-65 |
| | *FAP* | 2.2442 | n.s. |
| | *DCN* | 4.1488 | n.s. |

* Positive log2-fold changes represent increased gene expression in FdiAs and negative log2-fold changes represent increased gene expression in fibroblasts.
### Padj: p-values were corrected for FDR using the Benjamini-Hochberg method in R.

**Table S2.** Cell Lines used in this study

| Sl. No. | Cell Line # | Cell Type | Age (years) | Sex | Category |
|---|---|---|---|---|---|
| 1 | AG04455 | Fibroblast | 72 | Male | Aged Healthy Adult |
| 2 | AG09173* | Fibroblast | 75 | Female | Aged Healthy Adult |
| 3 | GM04506 | Fibroblast | 20 | Female | Young Healthy Adult |
| 4 | AG27611* | iPSC | 75 | Female | Aged Healthy Adult |

* Fibroblast and iPSC lines derived from the same individual.

#### SUPPLEMENTARY EXPERIMENTAL PROCEDURES

#### METHOD DETAILS

##### Immunocytochemistry

Cells were plated on glass coverslips for imaging. Cells were fixed with 4% PFA for 10 min at room temperature and washed thrice with PBST, followed by a 1-hr block/permeabilization in PBS containing 3% BSA (Sigma) and 0.1% Triton X-100 (Sigma). Primary antibodies were incubated in blocking solution overnight at 4°C. After PBST washes, secondary antibodies were incubated for 1 hour at room temperature in blocking solution, followed by PBST washes and nuclear staining with DAPI (1/10,000; Sigma-Aldrich). After washing, coverslips in PBS were imaged using a Leica DMi8 inverted fluorescence microscope or using a Zeiss LSM 780 upright confocal microscope. Primary and secondary antibodies used in the study are listed in the key resources table. All data for one experiment were acquired from cells cultured and processed in parallel and imaged using the same microscope settings.

**Image Analysis**

Images were acquired with the Leica DMi8 inverted fluorescence microscope, using LAS-X software, at 20X magnification (unless otherwise specified). 16-bit images were acquired, and exposure times were set to intensities as viewed under the microscope and were kept constant for each staining experiment. Confocal images were acquired with the Zeiss LSM 780 upright microscope using ZEN software. Images were acquired at 20x, 40x or 63x, at 1AU. Laser power for each wavelength was maintained throughout experiments. Secondary-only antibody controls were acquired using the same settings for all acquisitions. To quantify the number of GFAP and S100β positive cells, as well as other astrocyte markers, 2-3 stained coverslips from as many independent replicates were imaged per cell line, and cells were counted in 12-15 randomly selected fields of view. Acquired images were processed in the Fiji distribution of ImageJ (Schindelin et al., 2012). Total number of immunoreactive cells and the number of nuclei per field of view were counted in background subtracted (rolling ball radius: 50) images. For quantification of fluorescence intensities, three independent replicates were used, and 10-12 random fields of view were imaged. Background subtraction was carried out as before, and 120-150 ROIs were marked manually using corresponding phase contrast images. Absolute fluorescence intensities were then recorded per ROI, and mean intensities were represented.

**Whole-genome mRNA Sequencing (RNA-seq) and Analysis**

Total RNA was extracted from triplicate samples of fibroblasts (0 DIV) and FdiAs at 40 DIV using the All-Prep DNA/RNA mini kit (Qiagen). RNA was quantified using Nanodrop (Thermo Fisher Scientific), integrity assessed using Agilent Bioanalyzer and sequencing conducted by Novogene Co., Ltd. (UC Davis, CA). Paired-end 150 bp sequences were generated using the Illumina Novoseq 6000 platform, and raw fastq files were obtained. Raw files were checked for quality and pre-processed using the *fastp* algorithm (Chen et al., 2018). Reads were mapped to the Human Genome (hg38 version, UCSC genome database) using *Hisat2* (Kim et al., 2019), and raw counts were generated using *featureCounts* (Liao et al., 2014). Differential expression analyses were performed in *DESeq2* (Love et al., 2014), and cutoffs were set for log2FC > |1.5|. Statistical values were corrected for false discovery rates (FDR) using the Benjamini-Hochberg method implemented in R, and significance was set at FDR <0.05. Volcano plots were generated using the *EnhancedVolcano* package in R (Blighe et al., 2019). Gene set enrichment analysis (GSEA) for either GO Biological Processes or GO Molecular Function, or both, and KEGG pathways were performed using the *clusterProfiler* package in R (Wu et al., 2021; Yu et al., 2012), and corresponding dot plots were generated using the *DOSE* package (Yu et al., 2015). For comparison with previous astrocyte and other cell-type specific bulk RNA-Seq datasets (GSE73721, GSE84826, GSE97619, GSE97904, GSE101913, GSE104232, GSE168375, GSE174379, and GSE246073), raw fastq files were obtained, and analysis was performed as described above. Paired-end and single-end read data were processed separately, and count files generated from featureCounts were merged prior to DESeq2 analyses. All plots were generated using the *ggplot2* package in R. Principal component analysis and hierarchical clustering between our data, and other cell-type specific samples from the public data set were based on variance-stabilized transforms, generated in DESeq2, and plots were generated using *ggplot2* and *pheatmap* packages. For analysis of neuroinflammation phenotype data, vehicle-treated FdiAs and cytokine-stimulated FdiAs were run as triplicates, and analysis was conducted as detailed above. Differential expression was computed as before, GO and KEGG pathway analysis was conducted, and plots were generated.

Statistical tests for RNA-seq data from this study and other public datasets were performed using inbuilt normalization and significance correction tools in R (v4.2.2), featureCounts (v2.0.4), and DESeq2 (v 1.38.3). Adjusted p values (padj) were used to correct for multiple testing, and an FDR $<0.05$ was set for significance.

**Single-cell Sequencing (sc-RNA-seq) and Analysis**

Fibroblasts (Day 0), and FdiAs at different stages of conversion (Day 10, 30 and 50) were fixed and processed for sc-RNA-seq using a combinatorial indexing approach. Single-cell sequencing was carried out according to manufacturer's instructions using the Evercode Whole Transcriptome v2 Mini kit (Parse Biosciences, Seattle, USA). Sequencing was performed on an Illumina NovaSeq 6000 (UT Arlington), and fastq files were generated. Analysis was carried out using a cloud-based analysis platform, Trailmaker™ (https://app.trailmaker.parsebiosciences.com/). Briefly, sc-RNA-seq fastq files were uploaded for genomic alignment and quality control (QC), yielding normalized count matrices and Seurat objects for downstream analyses. Standard QC thresholds were utilized in processing the data, including ones based on the cellular distribution of unique molecular identifier (UMI) counts, mitochondrial content, linear relationship of the number of genes expressed and the number of UMIs per cell, and doublet identification. Using the R toolkit Seurat, Seurat objects were generated from the processed and log-normalized data. Data integration was achieved using the Harmony method, and non-linear dimensional reduction was performed using UMAP (minimum cosine distance of 0.3; clustering based on the Leiden method).

**Glutamate Uptake Assay**

To assess the glutamate uptake capacity of our astrocytes, we either supplemented HBSS with 125μM glutamate (Sigma) or with no glutamate (negative control), and loaded fibroblasts (D0), and FdiAs (D40) for 4-6 hours at 37C, 5% $CO_2$. Post-incubation, supplied buffer solutions with or without glutamate were collected. For EAAT2 inhibitor experiments, cells in HBSS were first exposed to 100μM WAY-213613 (Tocris

Biosciences) for 30 minutes, followed by the addition of glutamate. We first quantified total proteins in each solution using a conventional BCA assay (Invitrogen). Then, we assessed the total amount of glutamate remaining in the solution using the Glutamate uptake assay kit (Sigma) per the manufacturer's instructions. Glutamate levels in each sample were subtracted to no-glutamate supplemented conditions, and total levels of glutamate present per ug of protein were determined by absorbance measurements in the Glomax microplate reader (Promega). The percentage of glutamate uptake was calculated based on the initial glutamate supplied and represented as Mean +/- standard error of mean (SEM).

**Calcium imaging**

Fibroblasts (D0) and FdiAs (D42), plated on Matrigel-coated glass coverslips were washed thrice with Kreb's-Ringer HEPES-buffered solution (KRH buffer: 140 mM NaCl, 2 mM CaCl2, 4 mM KCl, 1 mM MgCl2, 10 mM Glucose, and 10 mM HEPES/NaOH, pH 7.4), and loaded with 4μM Fluo-4 AM (Invitrogen) containing 0.02% Pluronic F-127 (Invitrogen) in the same buffer for 30 minutes at 37°C. After incubation, cells were washed again, and de-esterified in the KRH buffer for 15 minutes at RT and imaged using a gravity-flow perfusion system with a Leica DMi8 Inverted Fluorescence Microscope at 488 nm, using a 20x magnification. Time-lapse images were acquired using the LAS-X software. For ATP-evoked calcium responses, baseline recordings were obtained for the first 30s, and ATP at a final concentration of 100 μM was loaded onto the cells. Residual ATP was washed out at 2 minutes using KRH buffer, and images were recorded until return to baseline (270s). Time-lapse image analyses were conducted using the Fiji distribution of ImageJ (Schindelin et al., 2012). Briefly, ROIs were marked manually using phase-contrast images, and fluorescence intensities across 270s were measured. Background intensities were subtracted at each time point based on the average of three fluorescence intensity recordings in non-cell ROIs. Normalized fluorescence intensities, $DF/F_0$ were measured considering average of 20 lowest fluorescence intensities as baseline. Calcium response curves were plotted as Mean +/- SEM, using custom scripts in R or Microsoft excel. The number of responding cells was computed as those responding within Mean +3 standard deviations (SDs) of baseline measurements, and peak amplitudes were generated based on maximum fluorescence intensities during each trace. These measures were plotted as Mean +/- SEM for 3-5 independent replicates. Independent t-tests were conducted considering either equal (% responding cells), or unequal (Peak amplitude) variance to estimate the significance between fibroblasts and FdiAs.

**Generation of iPSC-derived Neurons and Neuron-Astrocyte Co-culture**

For the generation of iPSC-derived neurons, an iPSC cell line derived from the same individual (aged healthy female) was obtained from Coriell. iPSCs were differentiated into neural stem cells using a dual SMAD inhibition, and embryoid body formation protocol as before (Mukherjee et al., 2019), and converted into neurons. Briefly, neural stem cells were plated on poly-L-ornithine and Laminin coated dishes in neural expansion media, and were differentiated into forebrain neurons. Neuronal maturation was done using the BrainPhys Neuronal Medium (Stem Cell Technologies). When cells reached Day 30, they were either replated on Poly-L-ornithine/Laminin coated coverslips or plated on mature FdiAs.

For co-culture experiments, FdiAs were grown as before, and sorted for CD49f+/ACSA-2+ cells at Day 18 of conversion using a BD FACS Aria II cell sorter (BD Biosciences). Post-sort, FdiAs were replated on Matrigel-coated coverslips and allowed to mature as before until Day 40. iPSC-neurons were then plated on Day 40 FdiAs and cultured in 1:1 BrainPhys: Astrocyte maturation media for at least 1 week. iPSC-iNs and co-cultures were fixed and stained as before for MAP2 and SYN1, and images were acquired using the Zeiss LSM 780 confocal microscope.

**Neurite Length and Puncta Analysis**

Z-stack images obtained using the confocal microscope were processed as before, and images were analyzed for neurite length (using MAP2+ staining), using the Simple Neurite Tracer (SNT) plugin (Arshadi et al., 2021) in the Fiji version of ImageJ. For SYN1+ puncta analyses, background subtracted images were independently analyzed and the number of SYN1+ puncta per 50um of MAP2 neurites was estimated.

**Cytokine Stimulation Assay**

To evaluate the neuroinflammatory potential of FdiAs, cells at D40 of differentiation were stimulated with IL-1a (Peprotech), TNFa (Peprotech) and complement C1q (Sigma) in astrocyte maturation media for 24 hours at 37C, 5% CO2. Cells were collected by manual scraping, and total bulk RNA was extracted as before and sequenced (see RNA-sequencing and analysis section). Differentially expressed genes between vehicle-treated or stimulated FdiAs were assessed using DESeq2.
